# karyoploteR: an R/Bioconductor package to plot customizable linear genomes displaying arbitrary data

**DOI:** 10.1101/122838

**Authors:** Bernat Gel, Eduard Serra

## Abstract

**Motivation:** Data visualization is a crucial tool for data exploration, analysis and interpretation. For the visualization of genomic data there lacks a tool to create customizable non-circular plots of whole genomes from any species.

**Results:** We have developed karyoploteR, an R/Bioconductor package to create linear chromosomal representations of any genome with genomic annotations and experimental data plotted along them. Plot creation process is inspired in R base graphics, with a main function creating karyoplots with no data and multiple additional functions, including custom functions written by the end-user, adding data and other graphical elements. This approach allows the creation of highly customizable plots from arbitrary data with complete freedom on data positioning and representation.

**Availability:** karyoploteR is released under Artistic-2.0 License. Source code and documentation are freely available through Bioconductor (http://www.bioconductor.org/packages/karyoploteR)

## 1 Introduction

Data visualization is an important part of data analysis. It efficiently summarizes complex data, facilitates exploration and can reveal non-obvious patterns in the data. A natural representation for genomic data is positioned along the genome next to the ideograms of the different chromosomes. This type of representation is specially useful to identify the relation between different types of experimental data and genomic annotations. Various genomic visualization tools are available. Circos (Krzywinski *et al*., 2009) produces highly customizable high quality circular plots, as does it’s R counterpart RCircos (Zhang *et al*., 2013). There are other R packages capable of plotting whole genome diagrams such as: ggbio (Yin *et al*., 2012), based on the grammar of graphics that can produce different plot types including ideogram and karyogram plots; IdeoViz (Pai and Ren, 2014), to plot binned data along the genome either as lines or bars; or chromPlot (Oróstica and Verdugo, 2016), to plot up to four datasets given in a predefined format. In addition, the Bioconductor package Gviz (Hahne and Ivanek, 2016) is a powerful tool to create track based plots of diverse biological data but it does not produce plots of the whole genome. There is a lack of a tool to create non-circular whole genome plots, able to plot arbitrary data in any organism and with ample customization capabilities.

Here we present karyoploteR, an extendable and customizable R/Bioconductor package to plot genome ideograms and genomic data positioned along them. It’s inspired on the R base graphics, building plots with multiple successive calls to simple plotting functions.

## 2 Features

The interface of karyoploteR and the process to create a complete plot is very similar to that of base R graphics. We first create a simple or even empty plot with an initializing function and then add additional graphic elements with successive calls to other plotting functions. The first call creates and initializes the graphical device and returns a *karyoplot* object with all the information needed to add data to it. The *karyoplot* object contains a coordinate change function mapping genomic coordinates into plotting coordinates, which is used by all plotting functions. Plotting functions are classified into three groups: the ones adding non-data elements to the plot (titles, chromosome names, …) (*kpAdd*\* functions) and two data plotting groups, low-level functions (*kp*\*) and high-level functions (*kpPlot*\*). karyoploteR also takes some ideas from Circos, such as not defining fixed tracks but leaving complete freedom to the user with respect to data positioning using the *r0* and *r1* parameters.

**Fig. 1.**
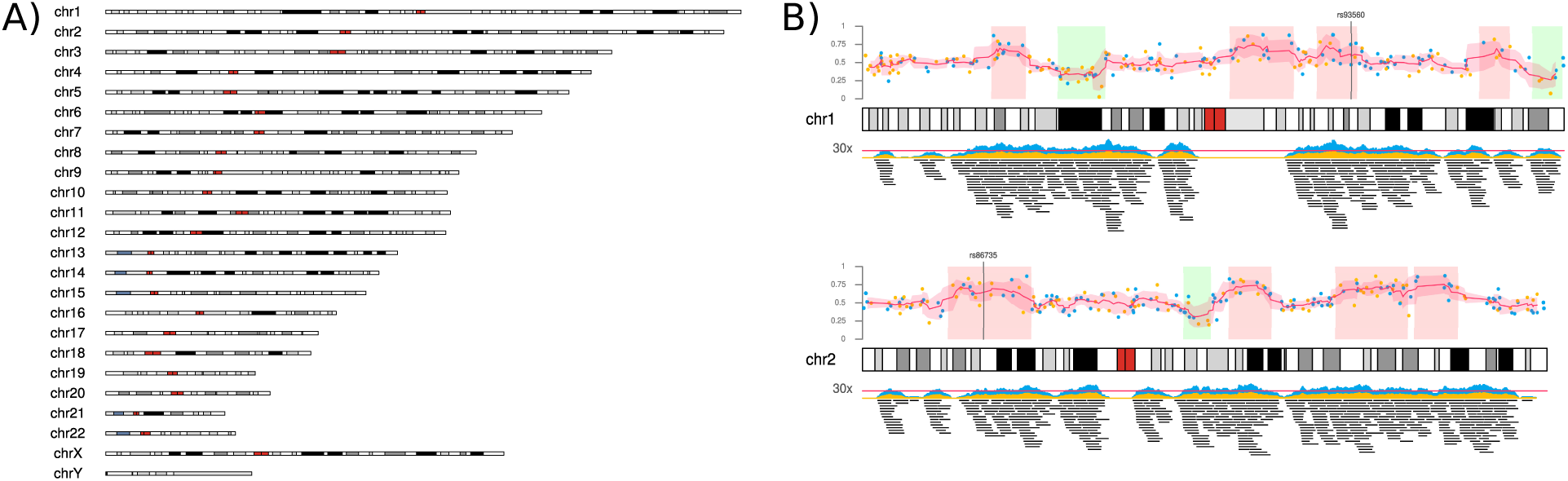
A) The complete human GRCh38 genome. This plot is created with the single command “plotKaryotype(genome=“hg38”)”. B) An example of a figure generated by karyoploteR representing different data types plotted in human chromosomes 1 and 2.

### 2.1 Ideogram plotting

Ideogram plotting is the basic functionality of karyoploteR. Default ideograms can be plotted with a single function call (Figure 1 A). However, it’s possible to customize them, positioning the chromosomes in different arrangements, representing just a subset of chromosomes or change whether the cytobands are included and how they are represented. It is also possible to create different data plotting regions either above or below the ideograms as well as customizing all sizings and margins by changing the values stored in *plot.params*. All elements in the karyoplot (main title, chromosome names, cytobands, …) are drawn by specific functions. These functions accept standard graphical parameters but it’s possible to swap them for custom functions if a larger level of customization is needed.

### 2.2 Not only human

karyoploteR is not restricted to human data in any way. It is possible to specify other organisms when creating a karyoplot. Genome data for a small set of organisms is included with the package and it will use functionality from regioneR Gel *et al*. (2016) to get it from UCSC or Bioconductor for other genomes. If an organism is not available anywhere, it is possible to plot it providing its genome information. Therefore, if required, it’s possible to create custom genomes for specific purposes.

### 2.3 Data plotting

Data plotting functions are divided in two groups: low-level and high-level. Low-level data plotting functions plot graphical primitives such as points, lines and polygons. Except for the additional *chr* parameter, they mimic the behaviour of their base graphics counterparts including the usage of most of the standard graphical parameters. These plotting functions offer a flexible signature and are completely data agnostic: they know nothing about biological concepts, giving the user total freedom on how to use them. High-level functions, in contrast, are used to create more complex data representations. They understand some basic concepts such as “genomic region” and they usually perform some kind of computation prior to *kpPlotRegions, kpPlotDensity* and *kpPlotBAMDensity* are examples of these type of functions.

### 2.4 Customization and extensibility

In addition to customizing sizings and margins and the using custom genomes, karyoploteR can be extended with custom plotting functions. All internal functions, including the main coordinate change function, are exported and documented in the package vignette. With this it is possible to create custom plotting functions adapted to specific data types and formats.

## 3 Conclusion

We have developed an R/Bioconductor package, karyoploteR, to plot arbitrary genomes with data positioned on them. It offers a flexible API inspired in R base graphics, with low-level functions to plot graphical primitives and high-level functions to plot complex data. The plots are highly customizable in data positioning and appearance and it is possible to extend the package functionality with custom plotting functions. More information and examples can be found at the package Bioconductor page and the GitHub repository https://github.com/bernatgel/karyoploter_examples.

## Acknowledgements

The authors thank Roberto Malinverni for his insightful comments and the IGTP HPC Core Facility and Iñaki Martínez de Ilarduya for his technical help.

## Funding

This work has been supported by: the Spanish Ministry of Science and Innovation, Carlos III Health Institute (ISCIII) (PI11/1609; PI14/00577)(RTICC RD12/0036/008) Plan Estatal de I + D + I 2013-2016, and co-financed by the FEDER program; the Government of Catalonia (2014 SGR 338); and the Spanish Association Against Cancer (AECC).

